# The genome of the endangered Dryas monkey provides new insights into the evolutionary history of the vervets

**DOI:** 10.1101/613273

**Authors:** T. van der Valk, C. M. Gonda, H. Silegowa, S. Almanza, I. Sifuentes-Romero, T. Hart, J. Hart, K. Detwiler, K. Guschanski

## Abstract

Genomic data can be a powerful tool for inferring ecology, behaviour and conservation needs of highly elusive species, particularly when other sources of information are hard to come by. Here we focus on the Dryas monkey (*Cercopithecus dryas*), an endangered primate endemic to the Congo Basin with cryptic behaviour and possibly less than 250 remaining individuals. Using whole genome sequencing data we show that the Dryas monkey represents a sister lineage to the vervets (*Chlorocebus sp.*) and has diverged from them around 1.4 million years ago with additional bi-directional gene flow ∼750,000 – ∼500,000 years ago that has likely involved the crossing of the Congo River. Together with evidence of gene flow across the Congo River in bonobos and okapis, our results suggest that the fluvial topology of the Congo River might have been more dynamic than previously recognised. Despite the presence of several homozygous loss-of-function mutations in genes associated with sperm mobility and immunity, we find high genetic diversity and low levels of inbreeding and genetic load in the studied Dryas monkey individual. This suggests that the current population carries sufficient genetic variability for long-term survival and might be larger than currently recognised. We thus provide an example of how genomic data can directly improve our understanding of highly elusive species.

## Introduction

The Dryas monkey (*Cercopithecus dryas*) is a little-known species of guenons endemic to the Congo Basin, previously only recorded from a single location (Figure 1). The recent discovery of a second geographically distinct population in the Lomami-Lualaba interfluve area has led to the elevation of its conservation status from critically endangered to endangered, however little is known about its population size, distribution range, behaviour, ecology and evolutionary history (Hart *et al.*, 2019). Based on pelage coloration, the Dryas monkey was first classified as the Central African representative of the Diana monkey (*Cercopithecus diana*) (Schwarz, 1932). However, later examinations of the few available specimens suggested that the Dryas monkey should be classified as a unique *Cercopithecus* species (Colyn, Gautier-Hion and van den Audenaerde, 1991). More recently, Guschanski *et al.* 2013 described the mitochondrial genome sequence of the Dryas monkey type specimen, preserved at the Royal Museum for Central Africa (Tervuren, Belgium), providing the first genetic data for this species. The mitochondrial genome-based phylogeny placed the Dryas monkey within the vervet genus *Chlorocebus*, supporting previously suggested grouping based on similarities in feeding behavior, locomotion and cranial morphology (Kuroda, Kano and Muhindo, 1985; Butynski, 2013). The vervets consist of six recognised species: Chl. *sabaeus, aethiops, tantalus, hilgerti, pygerythrus* and *cynosuros* (Zinner *et al.*, 2013). They are common in savannahs and riverine forests throughout sub-Saharan Africa, as well as on several Caribbean islands, where they were introduced during the colonial times (Figure 1A). However, the Dryas monkey is geographically isolated from all vervets and has numerous highly distinct morphological characteristics (Zinner *et al.*, 2013). As vervets are characterized by a dynamic demographic history with extensive hybridisation (Svardal *et al.*, 2017), including female-mediated gene flow, transfer of mitochondrial haplotypes between species can result in discordance between the mitochondrial tree and the true species evolutionary history. Thus, the phylogenetic placement of the Dryas monkey remains uncertain.

**Figure 1.**
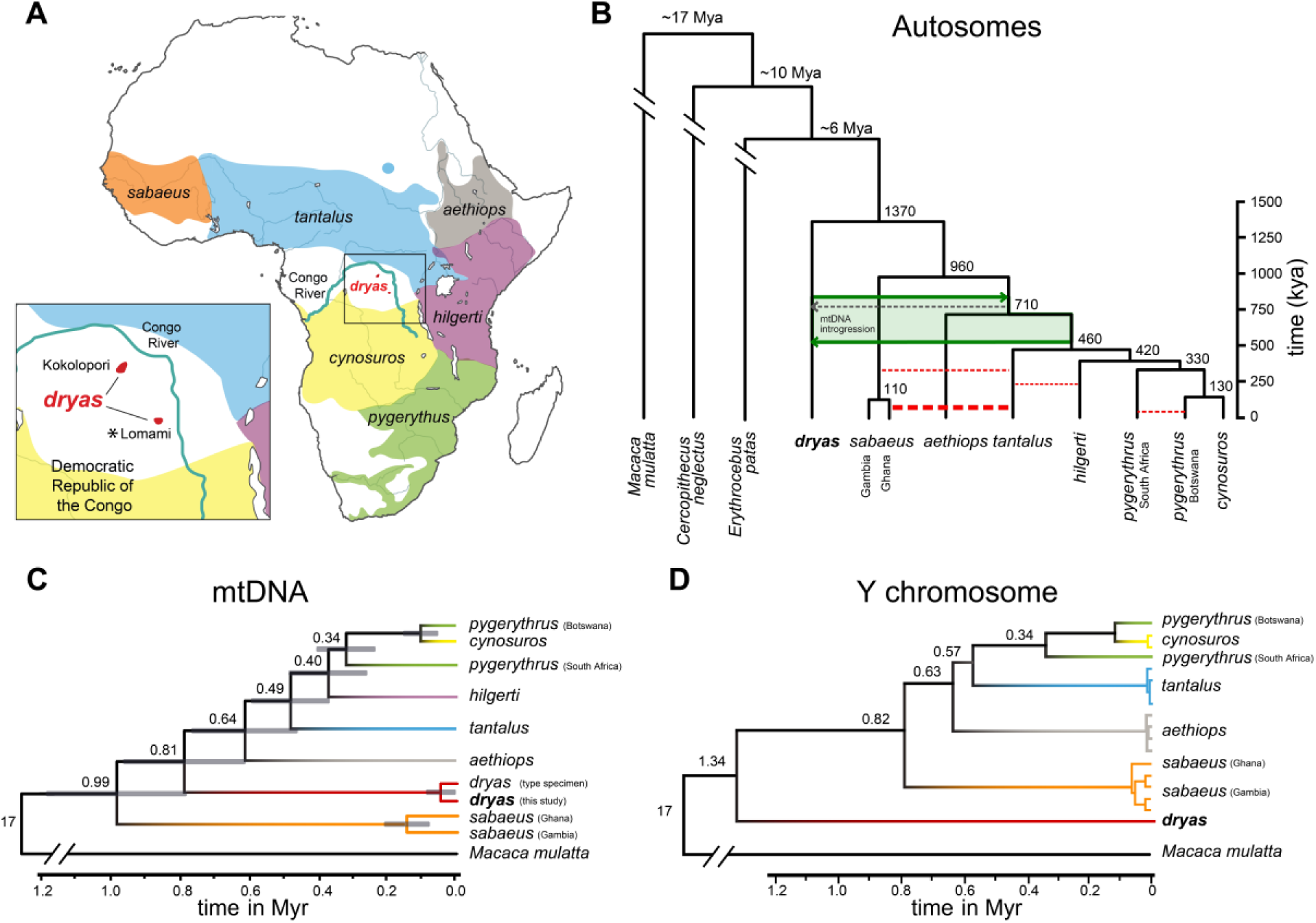
Phylogeny of the Dryas monkey and the vervets. **(A)** Distribution ranges of vervets. *Cercopithecus dryas* is currently known from only two isolated populations: The Kokolopori-Wamba and the Lomami-Lualaba interfluve. The sequenced Dryas monkey individual was sampled from the Lomami population as indicated by the asterisk. **(B)** The autosomal consensus phylogeny and MCMC dating of the Dryas monkey and the vervets in comparison to two other guenon genomes, using the rhesus macaque (*Macaca mulatta*) as outgroup. The tree topology is supported by multiple analyses (see Methods, Figure S2, S3). Red dotted lines depict previously reported admixture events between the different vervet species (Svardal *et al.*, 2017), with the line width corresponding to the relative strength of admixture. Green arrows show the best-supported timing of admixture between the Dryas monkey and the vervets. *Chl. sabaeus* individuals from Ghana have a higher proportion of shared derived alleles with the Dryas monkey than *Chl.* s*abaeus* individuals from Gambia, which is the result of a strong secondary admixture pulse between *Chl. tantalus* and the Ghana *Chl. sabaeus* population (Svardal *et al.*, 2017). Numbers at the nodes represent estimated mean divergence times (see Figure S3). (**C**) Mitochondrial phylogeny and divergence times, provided as numbers at the nodes. Grey blocks show the 95% confidence intervals as obtained with MCMCTree. (**D**) UPGMA Y-chromosomal phylogeny, calibrated using the rhesus macaque Y-chromosome as outgroup. Number at the nodes represent estimated divergence times.

With only two known populations and possibly fewer than 250 individuals, the Dryas monkey is of significant conservation concern (Hart *et al.*, 2019). The goal of this study was thus twofold: First, reconstruct the evolutionary and demographic history of the Dryas monkey and second, assess the long-term genetic viability of this species. To this end, we sequenced the genome of a male Dryas monkey at high coverage (33X), which represents the first genome-wide information available for this cryptic and little-known species.

## 2.0 Methods

### 2.1 Tissue sample collection and whole genome sequencing

Three Dryas monkey tissue samples were obtained from individuals from the Lomami National Park buffer zone in the Democratic Republic of Congo (Figure. 1A) and exported to Florida Atlantic University, United States under approved country-specific permits, where DNA was extracted using the Qiaqen DNeasy Blood & Tissue kit, following the manufactures protocol. For library preparation and sequencing the DNA extracts of two individuals were sent to Uppsala University, Sweden. The Illumina TruSeq DNA PCR-free kit was used for standard library preparation and the sample showing the highest DNA extract concentration was sequenced on one lane of the Illumina HiseqX platform (2x 150bp). In addition, we obtained previously published FASTQ data for all currently recognised vervet species from mainland Africa (Warren *et al.*, 2015; Svardal *et al.*, 2017) and divided them into two subsets: 23 *Chl. sabaeus*, 16 *Chl. aethiops*, 11 *Chl. tantalus*, 6 *Chl. hilgerti*, 16 *Chl. cynosuros* and 51 *Chl. pygerythrus* individuals sequenced on the Hiseq2000 platform at 1.9 – 6.8 X coverage (the low-coverage dataset), and one individual of each vervet species sequenced on the Genome-Analyzer II platform at 7.4 – 9.8X coverage (the medium-coverage dataset). We also obtained publicly available de-novo assembled genomes of two additional guenon species, the patas monkey (*Erythrocebus patas*, GenBank: GCA_004027335.1) and the De Brazza’s monkey (*Cercopithecus neglectus*, GenBank: GCA_004027615.1), and previously published high coverage genomes (27X and 31X) of two rhesus macaques (*Macaca mulatta*) to be used as outgroup (Xue *et al.*, 2016).

### 2.2 Alignment, variant detection and filtering

All FASTQ data were adapter and quality trimmed using Trimmomatic on recommended settings (Bolger, Lohse and Usadel, 2014) and initially aligned against the *Chlorocebus sabaeus* reference genome (ChlSab1.1) (Warren *et al.*, 2015) using bwa-mem (Li, 2013) with default settings. The de-novo patas and De Brazza’s monkey assemblies were first converted into 100 base pair non-overlapping FASTQ-reads and then aligned as above. For some analyses we noted significant reference bias arising from the ingroup position of *Chlorocebus sabaeus* reference with respect to the set of species studied here. As a result, FASTQ reads showing an alternative allele to the reference obtained lower mapping scores than reads carrying the references allele. This bias increases with genetic distance to the reference, and thus to avoid potential errors in our inferences, we additionally mapped all samples against the closest available outgroup reference (*Macaca mulatta*, Mmul_10) as above. All our subsequent analyses are based on the mappings to *Macaca mulatta*, unless stated otherwise. Samtools was used to filter out reads below a mapping quality of 30 (phred-scale) (Li *et al.*, 2009). Next, reads were realigned around indels using GATK IndelRealigner (McKenna *et al.*, 2010; DePristo *et al.*, 2011) and duplicates marked using Picard2.10.3 (https://broadinstitute.github.io/picard/). After these filtering steps we obtained a genome-wide coverage of 33X for the Dryas monkey. Next, we called single nucleotide polymorphisms (SNPs) with GATK UnifiedCaller outputting all sites (McKenna *et al.*, 2010; DePristo *et al.*, 2011). Raw variant calls were then hard filtered following the GATK best practices (Van der Auwera *et al.*, 2013). Additionally, we removed all sites below quality 30 (phred-scale), those with more than three times average genome-wide coverage across the dataset, sites for which more than 75% of samples had a non-reference allele in a heterozygous state, indels and sites within highly repetitive regions as identified from the repeatmask-track for the Mmul_10 reference using vcftools (Danecek *et al.*, 2011).

### 2.3 Cytochrome B sequencing

We amplified the cytochrome B sequence (1,140 bp) of the mitochondrial genome for all three Dryas monkey samples in five overlapping fragments (Haus *et al.*, 2013). PCR reactions were run in 25 µl final volume containing 1U GoTaq G2 Green Master Mix, 0.33 µM of each primer (forward and reverse, Table S1), 2 µl of genomic DNA and 6.5 µl ddH_2_0. The cycling conditions followed the procedure recommended by Haus et al. 2013 with a few modifications to the annealing temperatures of each primer, 94 °C for 2 minutes, followed by 40 cycles of 94 °C for 1 minute, primer-specific annealing temperature (Table S1) for 1 minute, 72 °C for 1 minute, and 72 °C for 5 minutes. All PCR products were checked on 2% agarose gels and then cleaned with the Wizard SV Gel and PCR Clean-up System from Promega (Madison, Wisconsin) and sent off for Sanger-sequencing along with the amplification primers to Molecular Cloning Laboratories (San Francisco, California). Sequence chromatograms were inspected by eye for accurate base calls and assembled using Geneious R11 11.0.5.

### 2.4 Autosomal phylogeny and divergence dating

First we used the vervet and the Dryas monkey SNP-calls to assess the genetic similarity between all individuals by applying a clustering algorithm. The R package *ape* was used to calculate pairwise distances among all individuals using the Tamura and Nei 1993 model, which allows for different rates of transitions and transversions, heterogeneous base frequencies, and between-site variation of the substitution rate (Tamura and Nei, 1993; Paradis, Claude and Strimmer, 2004). We then used the R package *phangorn* to construct a UPGMA phylogeny from the resulting pairwise distance matrix (Schliep, 2011).

Next, we reconstructed the phylogeny of the Dryas monkey in relation to the De Brazza’s monkey, patas monkey and the vervets using genome-wide phylogenetic methods. First, we generated haploidized sequences for one individual per species using ANGSD, by randomly selecting a single high quality base call (BaseQuality ≥30, read MapQuality ≥30, max depth below 3 times the genome-wide coverage) at each site in the reference genome, excluding sex-chromosomes and sites within repetitive regions (Korneliussen, Albrechtsen and Nielsen, 2014). We then concatenated autosomal sequences into a multi-species alignment file. Next, non-overlapping 350 kb genomic windows were extracted from the alignment and filtered for missing sites using PHAST v1.4 (Hubisz, Pollard and Siepel, 2011). After filtering, we excluded all windows with a length below 200 kb resulting in a final data set consisting of 3602 genomic windows (mean sequence length of 298 kb ±SD 18.5 kb). Individual approximately-maximum-likelihood gene trees were then generated for the set of windows using FastTree2 v2.1.10 (Price, Dehal and Arkin, 2010) and the GTR model of sequence evolution. Next, we constructed a coalescent species tree from the obtained gene trees with ASTRAL v5.6.2 (Zhang *et al.*, 2018) on default parameters. ASTRAL estimates branch length in coalescent units and uses local posterior probabilities to compute branch support for the species tree topology, which gives a more reliable measure of support than the standard multi-locus bootstrapping (Sayyari and Mirarab, 2016). Additionally, we obtained an extended majority rule consensus tree using the CONSENSE algorithm in PHYLIP v3.695 (Felsenstein, 2005). Finally, to explore phylogenetic conflict among the different gene trees, we created consensus networks with SplitsTree v4 using different threshold values (15% 20% and 25%) (Huson and Bryant, 2006).

After obtaining the consensus phylogeny, we used a Monte-Carlo-Markov-Chain (MCMC)-based method to estimate divergence times between the different species. For large alignments, calculation of the likelihood function during the MCMC is computationally expensive, and we therefore estimated divergence times using an approximate method as implemented in the software MCMC-Tree that significantly improves the speed of the MCMC calculations (Yang, 2007; Reis and Yang, 2011). For this analysis we used the independent molecular clock (the rates follow a log-normal distribution) and the JC69 evolutionary model with the following parameters as recommended by (Reis and Yang, 2011): alpha = 0, ncatG = 5, cleandata = 1, BDparas = 1 1 0.1, kappa_gamma = 6 2, alpha_gamma = 1 1, rgene_gamma = 2 20 1, sigma2_gamma = 1 10 1, finetune = 1: .1 .1 .1 .1 .1 .1, print = 1, burnin = 2000, sampfreq = 10, nsample = 20000. The divergence times were calibrated using a soft bound of 14.5– 19.6 Mya for the most recent common ancestor between the rhesus macaque (tribe *Papionini*) and the guenons (tribe *Cercopithecini*), as recently estimated in (Reis *et al.*, 2018). We repeated the MCMC run twice and checked for run convergence in R (R^2^ = 0.991).

We also estimated the divergence times between the Dryas monkey and the vervets using Pairwise Sequential Markovian Coalescent (PSMC) modelling specifically suited for low-coverage genomes (Cahill *et al.*, 2016). PSMC plots on artificial hybrid genomes (hPSMC) show a rapid increase in effective population size at the time when the two parental lineages diverge (Cahill *et al.*, 2016). Haploidized genomic sequences of the Dryas monkey and the vervets were merged using seqtk mergefa (https://github.com/lh3/seqtk.git) and we then ran hPSMC on artificial hybrids of all pairwise combination between Dryas monkey and the vervets (using four different vervet genomes per species) assuming a generation time of 8.5 years (Warren *et al.*, 2015) and a mutation rate per generation of 0.94×10^-8^ (Pfeifer, 2017).

### 2.5 Mitochondrial phylogeny

We *de-novo* assembled the mitochondrial genomes for the Dryas monkey, the vervets, and rhesus macaque with NOVOplasty (Dierckxsens, Mardulyn and Smits, 2016) with recommended settings, using a K-mer size of 39 and the *Chlorocebus sabaeus* (ChlSab1.1) mitochondrial reference genome as seed sequence. We also included the previously published mitochondrial genome sequence of the Dryas monkey type specimen (Guschanski *et al.*, 2013). Mitochondrial genomes were aligned using clustal omega with default settings (Sievers *et al.*, 2011). We then obtained a maximum-likelihood phylogeny in MEGAX, using the Tamura-Nei model with uniform rates among sites and running 1000 bootstrap replicates (Kumar *et al.*, 2018). Next we partitioned the protein-coding genes in the alignment into 1^st^,2^nd^ and 3^rd^ coding position, using the ChlSab1.1 annotation in Geneious 10.1.2 (https://www.geneious.com). We estimated divergence times with MCMC-tree (which allows for different evolutionary rates of the partitions) using the same parameters as for the autosomal divergence estimates, however with the likelihood (instead of approximate likelihood) function. Finally, we also aligned the cytochrome B sequences of two additional Dryas monkey individuals to all other mitochondrial genomes and reran the maximum likelihood estimates as above, this time only for the cytochrome-B sequence region.

### 2.6 Y-chromosome phylogeny

The mammalian Y chromosome sequence is enriched for repeats and palindromes, and thus accurate assembly from short-read data is challenging (Tomaszkiewicz, Medvedev and Makova, 2017; Kuderna *et al.*, 2019). We therefore obtained partial Y-chromosome consensus sequences using the filtered SNP calls. First, we identified all male individuals in our low-coverage dataset using the ratio of X-chromosome to autosomal coverage (Figure S1). Next, GATK FastaAlternateReferenceMaker was used to obtain a Y-chromosome consensus sequence for each male individual using the filtered variant calls as input. We masked all sites for which at least one individual showed a heterozygous call, as these represent SNP-calling errors. Additionally, we masked all repetitive regions and all sites for which one or more female individuals also showed a variant call, as these regions are likely enriched for SNP-errors due to mismappings. Given the scarcity of the retained genomic data (only 4% of the Y-chromosome was retained), we could not use any model-based phylogenetic approaches and instead constructed a UPGMA tree, as was done for the autosomal clustering, only including the filtered sites called in all male individuals. The tree was time calibrated using relative divergence to the rhesus Y chromosome, and a rhesus macaque - guenon divergence time of 17 million years (Reis *et al.*, 2018), assuming a uniform mutation rate.

### 2.7 Gene flow

We performed a model-free test of unbalanced allele sharing between the vervet individuals and the Dryas monkey (D-statistic) (Green *et al.*, 2010) using all autosomal bi-allelic sites called in all individuals. First we calculated D-statistics for all individual combinations as H1,H2,H3,H4, with the Dryas monkey as H3 and the rhesus macaque as the representative of the ancestral variant H4 (we excluded sites for which the two rhesus-macaque genomes were not identical). Next, we calculated frequency-stratified D-statistics by grouping all individuals by species and calculating the frequency of shared derived alleles within each species (e.g. for each genomic site where the Dryas monkey carries the derived allele, we calculated the frequency of the derived allele in each of the vervet species), again using the Dryas monkey genome as H3 and rhesus as H4. Genome-wide D-statistic on population level were then calculated for four different bins of derived allele frequencies (<0.25, 0.25-0.5, 0.5-0.75 and >0.75). A model-based estimate of introgression was obtained by constructing a maximum likelihood (ML) tree using TreeMix v. 1.12 (Pickrell and Pritchard, 2012), accounting for linkage disequilibrium (LD) by grouping sites in blocks of 1,000 SNPs (-k 1000). Based on the previous phylogenetic inferences, the Dryas monkey was set as root. Standard errors (-se) and bootstrap replicates (-bootstrap) were used to evaluate the confidence in the inferred tree topology and the weight of migration events. After constructing a maximum likelihood tree, migration events were added (-m) and iterated 50 times for each value of ‘m’ (1-10) to check for convergence in terms of the likelihood of the model as well as the explained variance following each addition of a migration event. The inferred maximum likelihood trees were visualised with the in-build TreeMix R script plotting functions.

To identify putatively introgressed regions in all vervet individuals, we performed a screen for such segments following a strategy outlined in (Martin, Davey and Jiggins, 2015). As this method is sensitive to the number of informative sites within windows, we used the mappings against the *Chlorocebus sabaeus* reference to maximise the number of informative sites (at the cost of reference bias). Briefly, in sliding windows of 10 kb we calculated 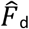 statistics (which is related to D-statistic but not subject to the same biases as D when calculated in sliding windows, (Martin, Davey and Jiggins, 2015)) and Dxy using all *Chl. sabaeus* individuals from Gambia as ingroup (H1) (as these showed the least amount of shared derived alleles with the Dryas monkey, see below). Genetic distance (Dxy) was calculated for each window to the Dryas monkey (Dxy_dryas-X_), and to *Chl. sabaeus* (Dxy_sabaeus(Gambia)-X_), where x refers to the focal individual. Next we calculated the ratio between Dxy_dryas-X_ / Dxy_sabaeus(Gambia)-X_ for each window. As Dxy is a measure of sequence divergence, introgressed windows between the Dryas monkey and non-*sabaeus* vervets are expected to have a relatively low Dxy_dryas-X_ and relatively high Dxy_sabaeus(Gambia)-X_. Next, windows showing an excess of shared derived alleles with the Dryas monkey (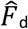 value > 2 times standard deviation) and an unusual low genetic divergence towards the Dryas monkey (the ratio of Dxy_Dryas-X_ / Dxy_sabaeus(Gambia)-X_ < the genome-wide average minus 2 times the standard deviation) were flagged as putatively introgressed. As regions with high 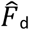 statistics tend to cluster in regions of low absolute divergence (*D*_*XY*_) (Martin, Davey and Jiggins, 2015), some of our identified introgressed regions might be false positives.

Additionally we used a topology weighting method, *Twisst*, for quantifying the relationships between the different taxa and visualizing how these relationships change across the genome (Martin and Van Belleghem, 2017). *Twisst* estimates the topology across the genome in sliding windows, where consecutive blocks of contrasting topologies to the majority topology are an indication of adaptive introgression. For this analysis, we first computationally phased the SNP-calls using shapeit (-phase– input-vcf beagle_out.vcf–window 0.1) (Delaneau *et al.*, 2013) and subsequently obtained topologies in sliding windows of 50 SNPs using phyml grouping of all samples by species, following the *Twisst* guidelines (https://github.com/simonhmartin/twisst.git). As the number of topologies increases exponentially with increasing number of species, running *Twisst* for many species becomes computationally unfeasible. We thus ran *Twisst* on the complete dataset of inferred phyml trees from the rhesus macaque, the Dryas monkey, *Chl. sabaeus, aethiops* and *cynosuros* samples (excluding *Chl. pygerythrus, hilgerti* and *tantalus* as we did not find strong differences in introgression proportions between these and *Chl. cynosuros*) using the complete method.

### 2.8 Selection and gene ontology enrichment of introgressed genes

Using the *Chlorocebus sabaeus* genome annotation (Warren *et al.*, 2015) we obtained for each individual all genes within putatively introgressed windows as identified from 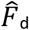 and Dxy statistics. A gene ontology enrichment was run for all putatively introgressed genes fixed in all non-*sabaeus* individuals in Blast2GO using Fisher’s exact test (Gotz *et al.*, 2008). Next, for all genes we obtained previously calculated selection score coefficients (XP-CLR selection scores) from Svardal *et al.* (2017). These selection scores are calculated for the different vervet populations and are based on a multilocus test of allele frequency differentiation, identifying regions in the genome where the change in allele frequency at the locus within the vervets occurred too quickly to be explained by drift using the *XP-CLR* method from (Chen, Patterson and Reich, 2010). Candidate genes for adaptive introgression were then identified as those with high gene-selection scores (top 10% of XP-CLR scores).

### 2.9 Demographic history

To infer long-term demographic history of the studied species, we used a pairwise sequentially Markovian coalescent model (PSMC) (Li and Durbin, 2011). As PSMC analysis is sensitive to the number of present genomic sites (Nadachowska-Brzyska *et al.*, 2016), we used the medium coverage vervets (7.4-9.8X coverage) and the Dryas monkey genome mapped to the *Chlorocebus sabeus* reference. We excluded sex chromosomes, repetitive regions and all sites for which read depth was less than five and higher than two times the genome-wide average. We scaled the PSMC output using a generation time of 8.5 years (Warren *et al.*, 2015) and a mutation rate of 0.94×10^-8^ per site per generation (Pfeifer, 2017), as above. Bootstrap replicates (n=100) were performed for the high-coverage Dryas monkey genome by splitting all chromosomal sequences into smaller segments using the splitfa implementation in the PSMC software and then randomly sampling with replacement from these fragments (Li and Durbin, 2011). Our Dryas monkey genome coverage (33X) differed strongly from that of the medium-coverage vervets. As limited coverage is known to biases PSMC results (Nadachowska-Brzyska *et al.*, 2016), we down-sampled the Dryas monkey genome to a similar coverage (8X) as the medium-coverage vervet genomes by randomly removing 75% of reads (samtools view –s 0.25) and repeated the PSMC analysis, allowing for relative comparisons between species. Demographic estimates from PSMC can also be biased by admixture events between divergent populations, giving a false signal of population size change (Hawks, 2017). We thus removed all putatively introgressed regions (see above) from the Dryas monkey genome and re-ran the PSMC analysis.

### 2.10 Heterozygosity and inbreeding

We measured genome-wide autosomal heterozygosity for all individuals with average genome coverage > 3X using realSFS as implemented in ANGSD, considering only uniquely mapping reads (-uniqueOnly 1) and bases with quality score above 19 (-minQ 20) (Fumagalli *et al.*, 2013; Korneliussen, Albrechtsen and Nielsen, 2014). ANGSD uses genotype-likelihoods, rather than variant calls, allowing for the incorporation of statistical uncertainty in low-coverage data and shows high accuracy in estimating heterozygosity for genomes above 3X coverage (Fumagalli, 2013; van der Valk *et al.*, 2019). Next, we used PLINK1.9 (Purcell *et al.*, 2007) to identify stretches of the genome in complete homozygosity (runs of homozygosity: ROH) for all individuals with average genome coverage > 3X. To this end, we ran sliding windows of 50 SNPs on the VCF files of all included genomes, requiring at least one SNP per 50kb. In each individual genome, we allowed for a maximum of one heterozygous and five missing calls per window before we considered the ROH to be broken.

### 2.11 Genetic load

To estimate genetic load in the Dryas monkey and vervets we used the mappings and genome annotation of the *Chlorocebus sabaeus*. The variant effect predictor tool (McLaren *et al.*, 2016) was used to identify loss-of-function mutations (transcript ablation, splice donor variant, splice acceptor variant, stop gained, frameshift variant, inframe insertion, inframe deletion, and splice region variant), missense and synonymous mutations on the filtered SNP calls. As an indication of mutational load, for each individual we counted the number of genes containing one or more loss-of-function and the total number of missense mutations divided by the number of synonymous mutations (dN/dS) (Fay, Wyckoff and Wu, 2001). We excluded all missense mutations within genes containing a loss-of-function mutation, as these are expected to behave neutrally. Dividing by the number of synonymous mutations mitigates species-specific biases, such as mapping bias due to the fact that the reference genome was derived from a *Chl. sabaeus* individual, coverage differences, and mutation rate (Xue *et al.*, 2015).

## 3.0 Results and discussion

### 3.1 The dynamic demographic history of the Dryas monkey and the vervets

In this study we present the first genome sequence of the Dryas monkey, and show that it is a sister lineage to all vervets (Figure 1B, S2, S3). Multiple phylogenomic approaches (MSC, Maximum likelihood and Bayesian) unambiguously support the same tree topology, thus contradicting the suggested placement within the *Chlorocebus* genus as inferred from the mitochondrial data (Figure 1C, Guschanski *et al.*, 2013). After analysing 3602 gene trees from autosomal genomic windows, we obtained a multi-species coalescent tree with maximum support values (lpp=1.0) for all nodes (Figure S2A). Our topology is consistent with the vervet phylogeny previously reported by Warren *et al.* 2015. Although a majority-rule consensus tree (Figure S2B) and network analyses at different threshold values showed some poorly resolved nodes within the vervet clade (Figure S2C), the position of Dryas monkey as sister lineage to all vervets remains unambiguous (Figure S2). By using an approximate likelihood MCMC method on all autosomes we estimate that the Dryas monkey diverged from the vervets around 1.4 million years ago, whereas the estimates of divergence times to both the genus *Cercopithecus* as exemplified by the De Brazza’s monkey (∼10.2Mya) and the genus *Erythrocebus* represented by the patas monkey (∼5.8 Mya) are much older (Figure 1B, S3). Artificial hybrid PSMC (hPSMC) analysis, which is based on mutation rate rather than calibration nodes, suggests a divergence with ongoing gene-flow between the Dryas monkey and the vervets starting at ∼2Mya ago with the final separation of the two lineages around 1.5 Mya (Figure S5). Both the MCMC and hPSMC analyses suggest that the radiation within the vervets started ca. 960 thousand years ago (Figure 1B, S6). Although our inferred divergence times are sensitive to both node calibration used in the MCMC and the mutation rate used for the hPSMC, they unambiguously support the sister relationship between the Dryas monkey and the *Chlorocebus* genus. Based on our analyses with only two representatives of the genus *Cercopithecus* (the Dryas monkey and de Brazza’s monkey), this genus appears paraphyletic. More guenon genomes will be needed to disentangle their complex evolutionary history, which may call for reconsideration of the taxonomic classification of the Dryas monkey.

The divergence times within the vervets inferred by us are more ancient than reported by Warren *et al.* (2015). This is likely explained by the use of a more ancient date for the divergence between the rhesus macaque and vervets as calibration node (14.5-19.6 Mya in this study versus 12 Mya in Warren *et al.* 2015), the mapping against an outgroup (*Macaca mulatta*) in contrast to Warren et. al 2015 who used an ingroup reference (*Chlorocebus sabaeus*), which likely artificially increased the similarity to *Chl. sabaeus* thus lowering the estimated divergence time, and the used methods (MCMC and hPSMC in this study versus pairwise differences in Warren et. al 2015). The Y-chromosome-based phylogeny shows the same topology and similar divergence time estimates as the autosomal tree (Figure 1D). This stands in stark contrast to the inferences based on mitochondrial sequences, which show that all four Dryas monkey individuals with available mtDNA sequence data are nested within the vervet genus *Chlorocebus* (Figure 1C, S7, S8).

The discrepancies in tree topologies derived from genomic regions with different inheritance modes (autosomal and mitochondrial) and the known history of introgression among the vervets (Svardal *et al.*, 2017) suggest a possible role of gene flow in shaping the evolutionary history of the Dryas monkey. Therefore, we explored if ancient admixture events can resolve the observed phylogenetic discordance between the autosomal and mitochondrial data. We found that *Chl. sabaeus* individuals from Gambia share significantly fewer derived alleles with the Dryas monkey (D-statistic) than all other vervets (Figure 2A). As derived alleles should be approximately equally frequent in all species under the scenario of incomplete lineage sorting without additional gene flow (Green *et al.*, 2010), such a pattern strongly suggests that alleles were exchanged between the Dryas monkey and the vervets excluding *Chl. sabaeus* that has split off from the others ∼960 kya (Figure 1B). *Chl. aethiops* individuals share significantly fewer derived alleles with the Dryas monkey than *Chl. tantalus, hilgerti, cynosuros*, and *pygerythrus* (Figure 2A), suggesting that gene flow likely occurred for an extended period of time, at least until after the separation of *Chl. aethiops* from the common ancestor of the other vervets (∼710kya, Figure 1B). As *Chl. tantalus, hilgerti, cynosuros* and *pygerythrus* vervets share a similar amount of derived alleles with the Dryas monkey, gene flow most likely ended before this group radiated (∼460kya, Figure 1B). The small observed differences in the D-statistic between *Chl. tantalus, hilgerti, cynosuros* and *pygerythrus* (±12%, Figure 2A) could be the result of drift and selection due to population size differences among these vervet species or hint at ancient substructure within these species (Slatkin and Pollack, 2008; Svardal *et al.*, 2017). The inferred history of gene flow is concordant with the mitochondrial phylogeny, which suggests that the Dryas monkey mitochondrial genome was introgressed from the common ancestor of all non-*sabaeus* vervets ∼810 kya (Figure 1B and 1C). Gene flow between the common ancestor of the non-*sabaeus* vervets and the Dryas monkey is also supported by TreeMix analyses (Figure S9), but we note that these inferences have to be interpreted with caution, as this model-based approach relies on accurate allele frequency estimates, which are absent for the Dryas monkey population, as it is represented by a single individual.

**Figure 2.**
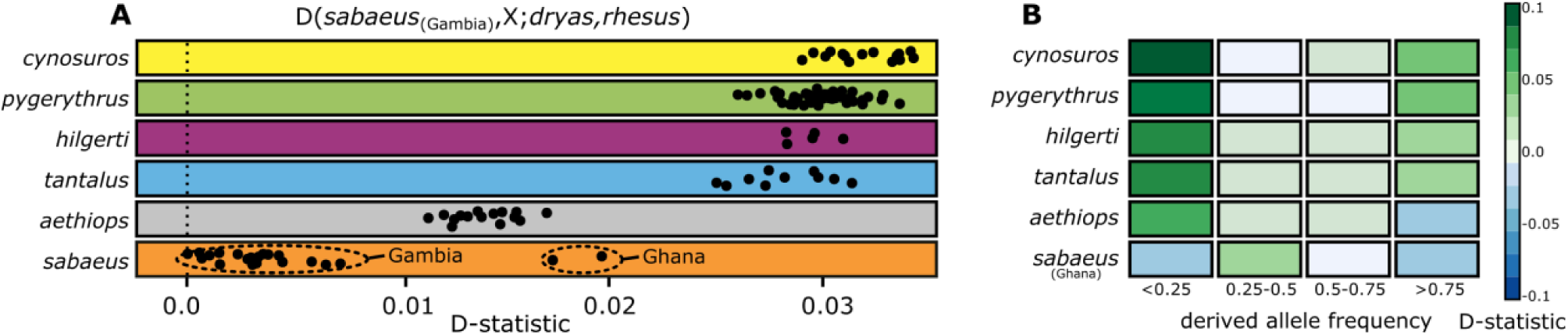
Genome-wide statistics support gene flow between the Dryas monkey and the vervets. (**A**) Obtained D-statistics for individual comparisons. Each dot depicts a comparison of a single vervet genome (X) with the *Chl. sabaeus* individual that showed the least amount of shared derived alleles with the Dryas monkey in all individual comparisons. (**B**) D-statistics stratified by derived allele frequency for each species, using all *Chl. sabaeus*_Gambia_ individuals as ingroup.

In contrast to the *Chl. sabaeus* individuals from Gambia, *Chl. sabaeus* vervets from Ghana also carry an excess of shared derived alleles with the Dryas monkey (Figure 2A). The Ghanese *Chl. sabaeus* population recently hybridised with *Chl. tantalus* and thus a large proportion of their genome (∼15%) is of recent *Chl. tantalus* ancestry (Figure 1B) (Svardal *et al.*, 2017). As *Chl. tantalus* individuals carry many shared derived alleles with the Dryas monkey, this secondary introgression event likely led to the introduction of Dryas monkey alleles into the Ghanese *Chl. sabaeus,* explaining the high D-statistics in this population.

Next, we obtained approximations of the directionality of gene flow using frequency-stratified D-statistics as in (de Manuel *et al.*, 2016). We found that the vervet populations carry derived alleles shared with the Dryas monkey at either low or high frequency, but few such alleles are found at intermediate frequencies (Figure 2B). High frequency alleles in the donor population are more likely to be introgressed and are subsequently present at low frequency in the recipient population (Kuhlwilm *et al.*, 2016). Therefore, our observation is consistent with bi-directional gene flow between the Dryas monkey and the non-*sabaeus* vervets. The general direction of the gene flow appears to have been dominated by the introgression from the Dryas monkey into the non-*sabaeus* vervets, as we observe an overall higher proportion of low-frequency shared derived alleles in the vervets (Figure 2B). This difference is particularly pronounced in *Chl. aethiops*, suggesting that the gene flow was primarily from the Dryas monkey into the vervets before *Chl. aethiops* separated from the common ancestor of *Chl. tantalus, hilgerti, cynosuros* and *pygerythrus*. After this split, gene flow likely became more bi-directional with increased proportion of introgression events into the Dryas monkey, as evidenced by the presence of high frequency putatively introgressed alleles in *Chl. tantalus, hilgerti, cynosuros* and *pygerythrus*. An alternative explanation is that the observed allele frequencies are driven by selection, as introgressed alleles might be on average selected against (Schumer *et al.*, 2018). It is also noteworthy that all four Dryas monkey individuals for which mitochondrial sequence data are available carry the vervet-like mitochondrial haplotype (Figure S8), which must have been introduced into the population through female-mediated gene flow. Given that the three Dryas monkeys in this study come from a different population than the Dryas monkey museum type specimen, this vervet-like haplotype is most likely fixed (or present at high frequency) in the *C. dryas* population.

The putatively introgressed alleles in the *Chl. sabaeus* population from Ghana are found at intermediate frequency (> 0.25 and < 0.50) (Figure 2B), which is in agreement with the indirect introduction of these alleles through recent introgression from *Chl. tantalus*. As the *Chl. tantalus* population carries derived alleles at high and low frequency (Figure 2B), the Ghanese *Chl. sabaeus* population received a mixture of both high and low frequency alleles, resulting in the observed intermediate frequency.

Using approaches that are relatively insensitive to demographic processes (e.g. genetic drift and changes in effective population size), we obtained strong support for the presence of gene flow between the Dryas monkey and the vervets, but we caution that they may incorrectly infer gene flow in situations with ancestral subdivision (Slatkin and Pollack, 2008). However, such ancestral population structure would have to persist over an extended period of time, encompassing multiple speciation events. Furthermore, our inferences of gene flow are supported by the discordance between the nuclear and mitochondrial phylogenies. Thus, gene flow seems to be the most parsimonious explanation.

### 3.2 Identifying introgressed regions and inferring their functional significance

Using D_xy_ and 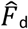 statistic (Martin, Davey and Jiggins, 2015) we identified putatively introgressed regions in sliding windows of 10.000 bp for each individual. As expected under the scenario of secondary gene flow, windows containing an excess of shared derived alleles with the Dryas monkey have low genetic divergence (D_xy_) to the Dryas monkey and high genetic divergence to *Chl. sabaeus* (Figure S10) (de Manuel *et al.*, 2016). Summing over all putatively introgressed windows, we roughly estimate that 0.4% - 0.9% in Gambia *Chl. sabaeus*, 1.6 – 2.4% in *Chl. aethiops* and 2.7% - 4.8% of the genome in the other vervets show as signature of introgression with the Dryas monkey. However we note that we can only identify the excess of Dryas monkey ancestry over that present in the *Chl. sabaeus* population. These estimates are thus likely a lower bound, as the *Chl. sabaeus* population might carry some Dryas monkey ancestry due to secondary gene-flow with other vervet species (Figure 1B, Svardal *et al.*, 2017). We estimate that putatively introgressed haplotypes average less than 10,000 bp and identify putatively introgressed blocks of up to 180 kb (Figure S11). However, we note that we cannot estimate the length of introgressed blocks below the length of 10,000 bp, as such short windows do not provide sufficient number of informative sites. The similar length of putatively introgressed haplotypes in all non-*sabaeus* vervet species strongly supports that gene flow occurred in their common ancestor. The putatively introgressed haplotypes into the Ghanese and (to a lesser extent) Gambian *Chl. sabaeus* populations were later introduced during secondary gene flow with *Chl. tantalus*, as an independent recent gene flow event from the Dryas monkey into these *Chl.* sabaeus individuals would have resulted in significantly longer haplotypes (Liang and Nielsen, 2014). However, we caution that our ability to accurately identify the haplotype lengths is low given the short length of the introgressed haplotypes and a single available Dryas monkey genome.

Interestingly, the proposed gene-flow between bonobos (*Pan paniscus*) and non-western chimpanzees (*Pan troglodytes*), which have overlapping distribution range with the Dryas monkey and the vervets, respectively, possibly occurred around a similar time period, (∼500.000 years ago, de Manuel *et al.*, 2016). It is noteworthy that the gene-flow that occurred between the Dryas monkey and the vervets most likely involved the crossing of the Congo River (Figure 1), previously thought to be an impenetrable barrier for mammals (Colyn, 1987; Colyn, Gautier-Hion and Verheyen, 1991; Colyn and Deleporte, 2004; Eriksson *et al.*, 2004). A similar scenario of cross-Congo River gene flow was proposed for the okapi populations, and bonobos-chimpanzees (Stanton *et al.*, 2014; de Manuel *et al.*, 2016), suggesting that the fluvial topology of the Congo River and the geology within the Congo basin might have been more dynamic 750,000 - 500,000 years ago than previously recognised (Beadle, 1981; Stankiewicz and de Wit, 2006).

Having identified introgressed regions, we explored if they may carry functional significance. First, we used *Twisst* to estimate the most likely topology along short sliding windows (50 SNPs), where multiple consecutive windows showing a contrasting topology to the majority topology can indicate adaptive introgression (Martin and Van Belleghem, 2017). Although this analysis supported our inferred species topology with additionally high support of introgression between the Dryas monkey and the vervets, we did not detect strong signals of long introgressed blocks, likely due to the very ancient timing of gene flow (Figure S11). As *Twisst* does not distinguish between incomplete lineage sorting and introgression for very short windows, we focused on introgressed regions identified with 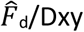 statistics. We find that genes previously identified to be under strong selection within vervets (top 10% of genes under the strongest selective pressure (Svardal *et al.*, 2017)) are less often introgressed with the Dryas monkey (average introgression frequency 0.021) than genes that did not experience strong selection (bottom 10%; average allele frequency 0.029). This may indicate weak selection against introgressed genes on average, which may be deleterious in the non-host background, a pattern also observed for Neanderthal genes in *Homo sapiens* and bonobo genes in the chimpanzee genetic background (Sankararaman *et al.*, 2014; Schumer *et al.*, 2018). However, we can not exclude that the difference in these gene frequencies is due to stochastic events or drift. To identify genes with adaptive functions, we focused on 109 putatively introgressed genes that are fixed in all non-*sabaeus* vervets. Gene ontology analysis did not reveal significant enrichment for any functional category (FDR=1). Vervets are the natural host of the simian immunodeficiency virus (SIV) and the genes under strongest selection in the vervets are related to immunity against this virus (Svardal *et al.*, 2017). We find POU2F1, AEBP2, and PDCD6IP among fixed putatively introgressed genes in all non-*sabaeus* individuals. POU2F1 is a member of the pathway involved in the formation of the HIV-1 elongation complex in humans (Sturm *et al.*, 1993), AEBP2 is a RNA polymerase II repressor (Kim, Kang and Kim, 2009) known to interact with viral transcription (Zhou and Rana, 2002; Debaisieux *et al.*, 2012), and PDCD6IP is involved in virus budding of the human immunodeficiency and other lentiviruses (Strack *et al.*, 2003; von Schwedler *et al.*, 2003). However, the SIV resistance related genes that experienced the strongest selection in vervets (e.g. RANBP3, NFIX, CD68, FXR2 and KDM6B) do not show a signal of introgression between vervets and the Dryas monkey. Therefore, while adaptive importance can be plausible for some of the introgressed loci, it does not appear to be a strong driver for retaining particular gene classes.

### 3.3 Genomic view on conservation of the endangered Dryas monkey

The Dryas monkey is considered the only representative of the Dryas species group (Grubb *et al.*, 2003) and is listed as endangered in the IUCN Red List due to its small population size of ca. 250 individuals (Hart *et al.*, 2019). We therefore used demographic modelling and genome-wide measures of heterozygosity and inbreeding to assess the long- and short-term population history of the Dryas monkey. Pairwise Sequential Markovian Coalescent (PSMC) analysis of the Dryas monkey genome revealed a dynamic evolutionary history, with a marked increase in effective population size starting ca. 600,000 years ago, followed by continuous decline in the last ∼150,000 years (Figure 3A). To eliminate the possibility that our PSMC inferences are driven by the increased heterozygosity due to gene flow, we removed all putatively introgressed regions and repeated the PSMC analysis, which produced a highly similar result (Figure 3A). We therefore suggest that the population size increase in the Dryas monkey and the associated likely range expansion facilitated secondary contact between the Dryas monkey and the vervets.

**Figure 3.**
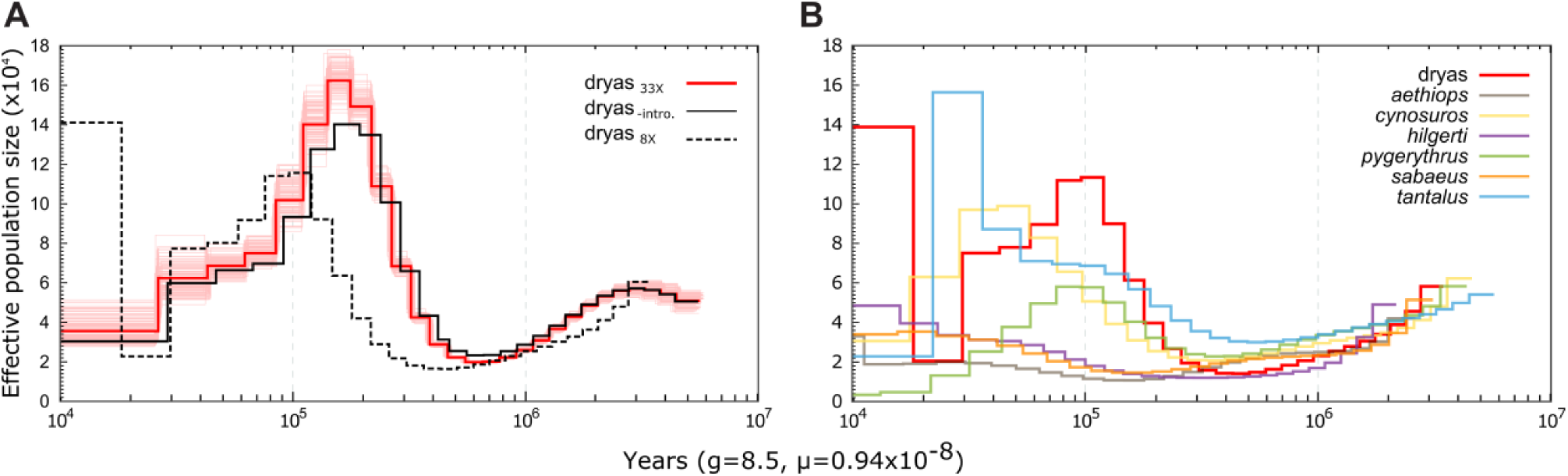
Demographic history of the Dryas monkey and the vervets. (**A**) PSMC analyses for the repeat-filtered Dryas monkey genome (red), after removing putatively introgressed regions (black), and after down-sampling to 8x coverage (dotted line). As the curve is strongly shifted at lower coverage, the down-sampled genome was used for between-species comparison in (B). (B) PSMC analysis of medium coverage genomes (7.4-9.8X) for all vervets and the down-sampled Dryas monkey genome (8X).

As previously reported, low genomic coverage shifts the PSMC trajectory and makes inference less reliable, particularly for more recent time periods (Nadachowska-Brzyska et al., 2016). Therefore, to allow for demographic comparisons to the vervets, we re-ran the PSMC on the Dryas monkey genome down-sampled to similar coverage as the genomic data available for the vervets. We indeed find a strong shift of the PSMC trajectory as a result of reduced coverage to more recent times and lower effective population size (Figure 3A). Nonetheless, bias related to coverage affect the different vervet genomes in a similar manner (Nadachowska-Brzyska et al., 2016), thus relative comparison between the species suggest that 100,000 – 300,000 years ago the Dryas monkey population was the largest among all vervets (Figure 3B).

The genetic diversity of the Dryas monkey (measured as between chromosome-pair differences), a proxy for the adaptive potential of a population (Lande and Shannon, 1996), is high compared to that of the much more abundant vervets (Figure 4A). The Dryas monkey individual also shows no signs of excessive recent inbreeding, which would manifest itself in a high fraction of the genome contained in long tracts of homozygosity (> 2.5 Mb) (Figure 4B). To estimate genetic load, we identified all genes in the Dryas monkey genome containing one or more loss-of-function (LoF) mutations and identified all missense mutations in genes other than those already containing LoF-mutation(s) (as such mutations likely behave neutrally). We find multiple genes in the Dryas monkey containing a homozygous LoF mutation associated with a disease phenotype in humans (n = 27), including in SEPT12, associated with reduced sperm mobility (Kuo *et al.*, 2015) and in SLAMF9, associated with reduced immunity to tapeworm infections (Cárdenas *et al.*, 2014). However, genome-wide measures of genetic load, measured as the ratio between LoF or missense and synonymous mutations (dN/dS), do not show an increased genomic burden of deleterious mutations in the Dryas monkey compared to the much more abundant and widely distributed vervets (Figure 4C-D). The demographic history and the genome-wide measures of genetic diversity and genetic load of the Dryas monkey thus suggest that the population of this endangered primate might be larger than currently recognised and has good chances for long-term survival, if appropriate conservation measures are implemented.

**Figure 4.**
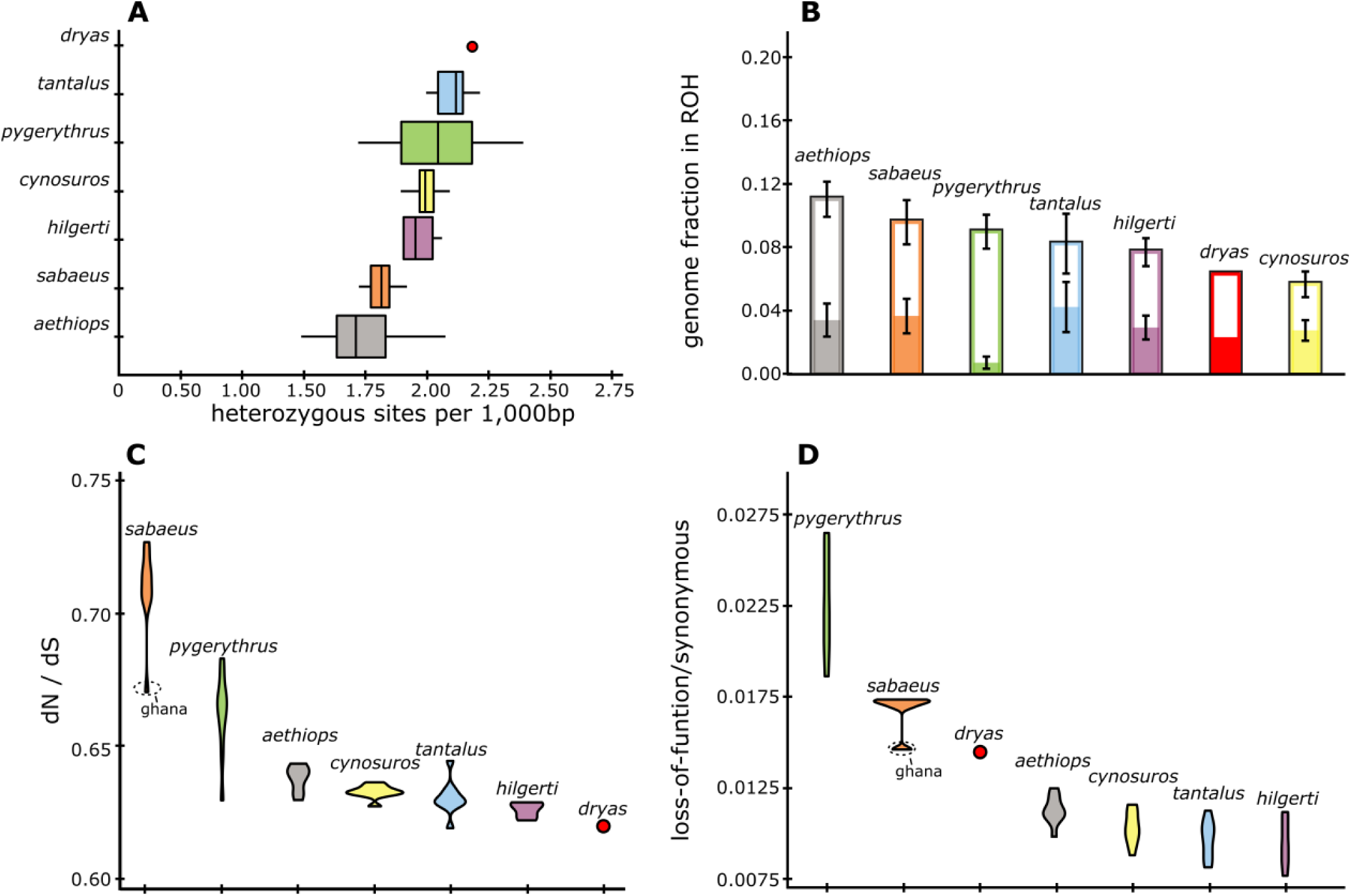
Genome-wide measures of genetic diversity and genetic load in the Dryas monkey. (**A**) Average number of heterozygous sites per 1,000 base pairs. (**B**) Fraction of the genome in runs of homozygosity (ROH) above 100 kb (open bars) and fraction of the genome in ROH > 2.5 Mb (solid bars). (**C**) Ratio of missense to synonymous mutations, excluding all missense mutations within genes containing one or more LOF mutations. (**D**) Ratio of LOF to synonymous mutations, counting genes with more than one LOF mutation only once.

## Acknowledgments

We thank the DRC government and ICCN for facilitating sample collection and the 200 Mammals Consortium for providing assemblies of *Cercopithecus neglectus* and *Erythrocebus patas*. Sequencing was performed by the SNP&SEQ Technology Platform in Uppsala. The facility is part of the National Genomics Infrastructure (NGI) Sweden and Science for Life Laboratory. The SNP&SEQ Platform is also supported by the Swedish Research Council and the Knut and Alice Wallenberg Foundation. The authors acknowledge support from the Uppsala Multidisciplinary Centre for Advanced Computational Science for assistance with massively parallel sequencing and access to the UPPMAX computational infrastructure. This work was supported by FORMAS (2016-00835) to K.G.

## Data accessibility

Whole genome sequence data generated in this study is available in the European nucleotide archive under accession number PRJEB32105. Cytochrome B sequences are available in the genbank under accession number XXX.

## Supplementary figures

**Table S1.**
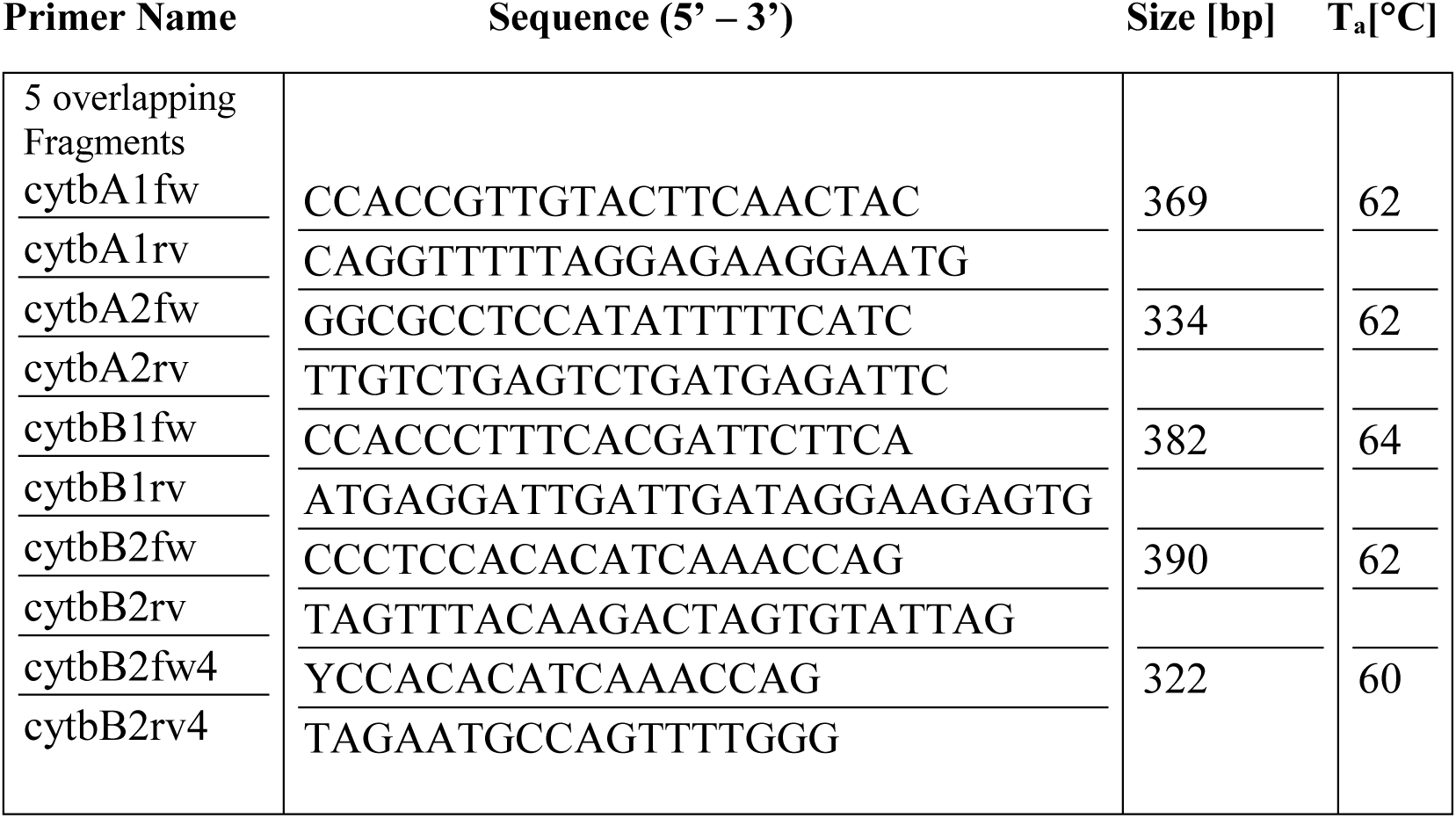
Primer sequences and annealing temperatures used for cytochrome B sequencing.

**Figure S1.**
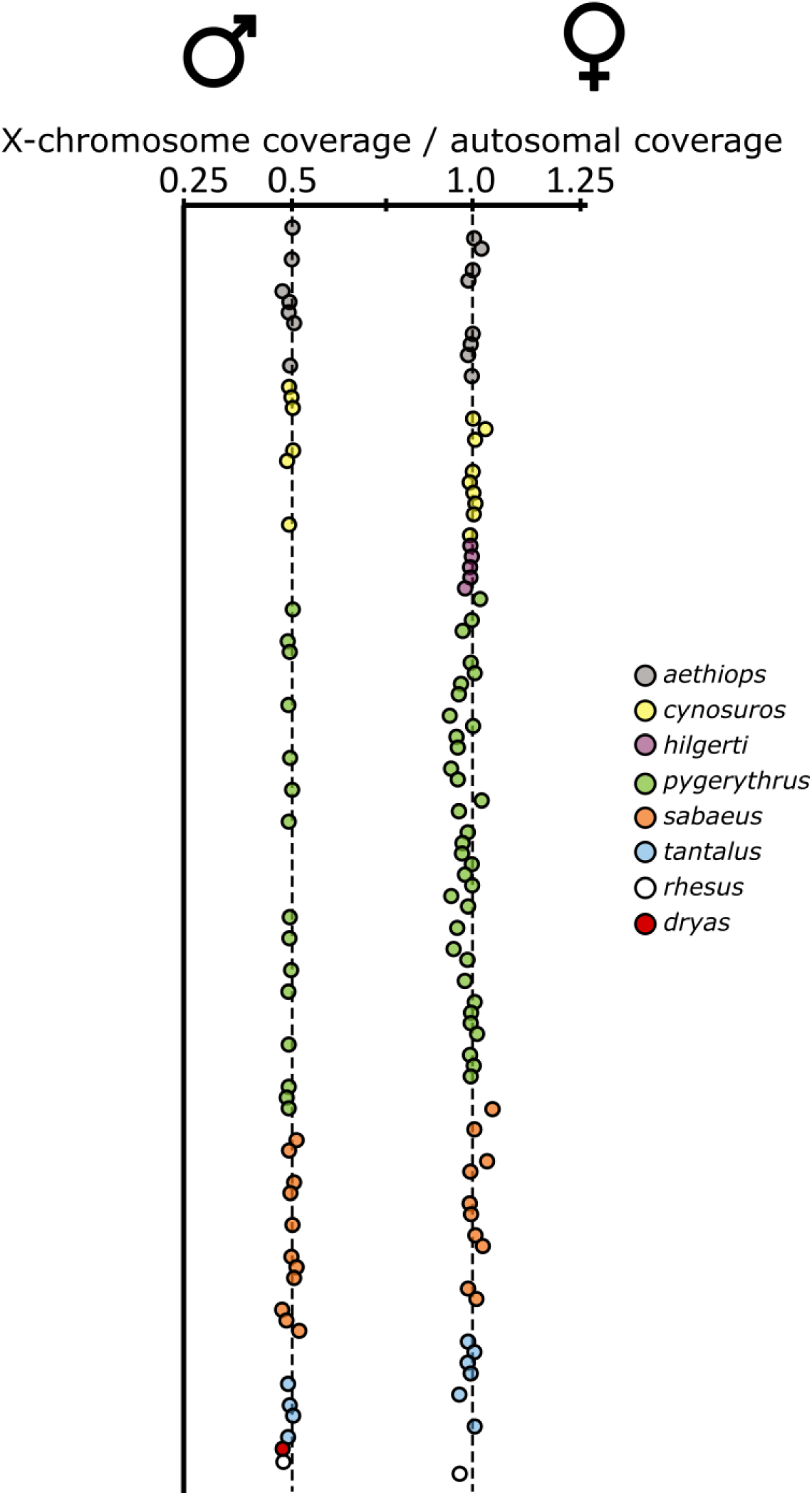
Sexing individuals based on the ratio of X-chromosomal to autosomal coverage.

**Figure S2.**
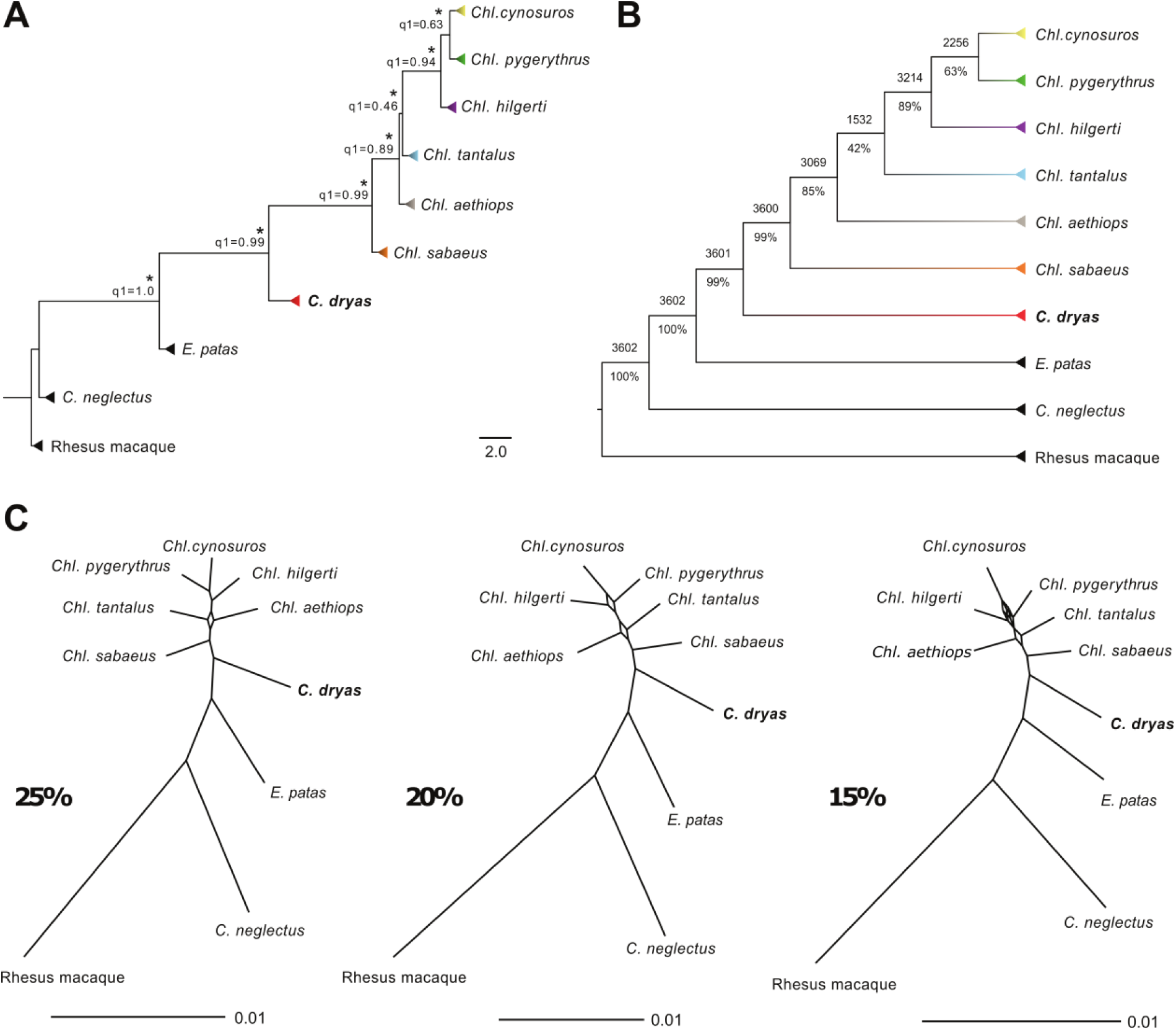
Phylogeny of the Dryas monkey and the vervets. **(A)** MSC-based species trees generated by ASTRAL using 3602 autosomal genomic windows. The tree was rooted with the rhesus macaque. Branch lengths are given in coalescent units and are an indicator of gene-tree discordance. The normalized quartet score of this topology is 0.87. Asterisk symbols at nodes indicate maximum local quartet support posterior probabilities (lpp=1.0) and *q1* values displayed at each node show the percentage of quartets in all gene trees that agree with that branch topology. (**B)** Majority-rule consensus tree obtained from 3602 autosomal gene-trees using CONSENSE in PHYLIP v3.695. The number above each branch shows the total number of trees out of all 3602 gene trees supporting the given branch and the number below corresponds to its percentage. **(C)** SplitsTree consensus networks using 3602 gene trees at median thresholds of 25%, 20% and 15%.

**Figure S3.**
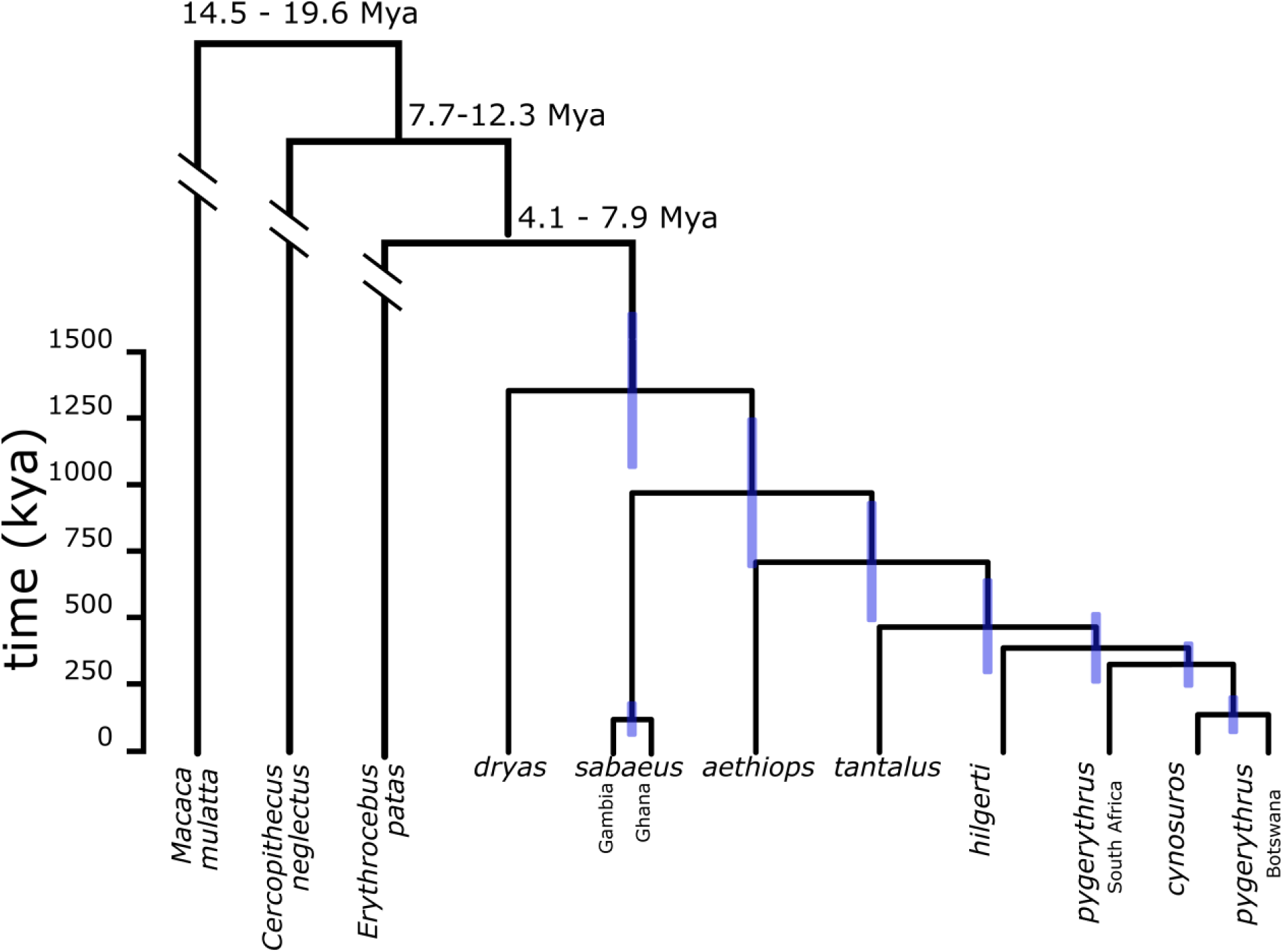
The autosomal consensus phylogeny and MCMC dating. Same as figure 1B but depicting 95% confidence intervals for the estimated divergence times. Confidence intervals for the oldest nodes are depicted with numbers for clarity.

**Figure S4.**
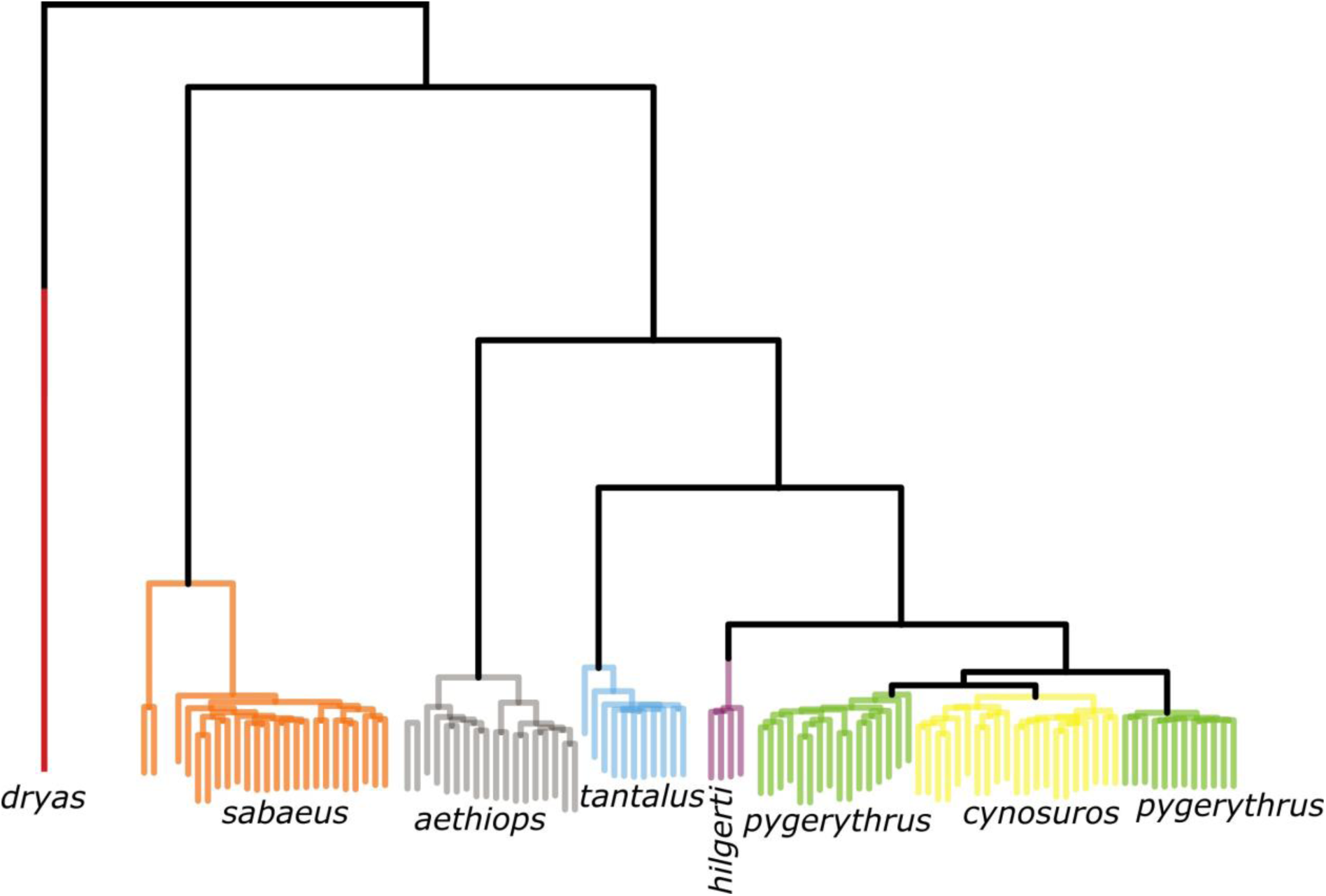
Evolutionary relationships of the Dryas monkey and the vervets. The tree shows the UPGMA clustering of all individuals used in this study based on all bi-allelic autosomal sites.

**Figure S5.**
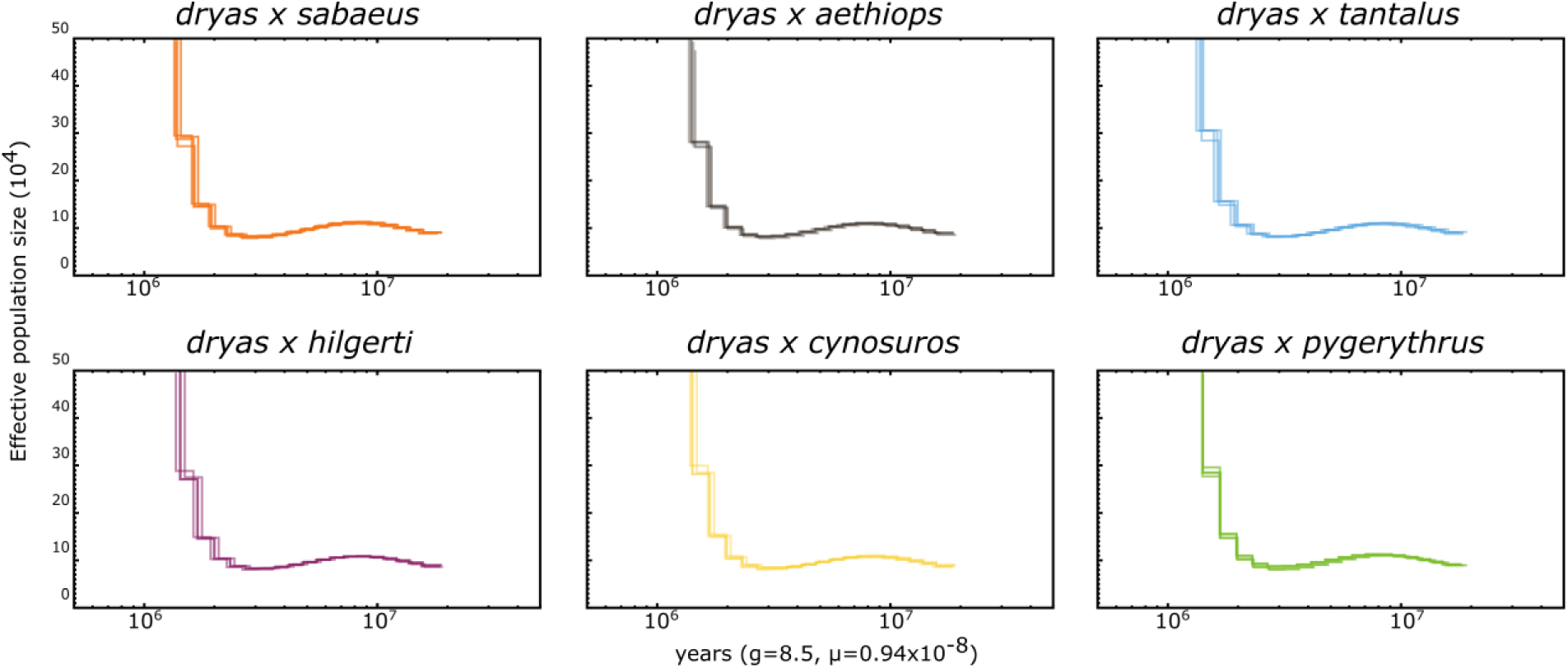
Artificial hybrid PSMC (hPSMC) of the Dryas monkey and the vervets. hPSMC was run on pairwise haploidized genomes for the Dryas monkey and four individuals from each vervet species assuming a generation time of 8.5 years and a mutation rate per generation of 0.94×10^-8^. PSMC plots on artificial hybrid genomes (hPSMC) show a rapid increase in effective population size at the time when the two parental lineages diverged. A stepwise increase in Ne is an indication of a population split with ongoing gene-flow. Note that the rapid increase in effective population size occurs at the same time for each combination of Dryas monkey and vervet species, suggesting that the Dryas monkey is equidistant to each of them.

**Figure S6.**
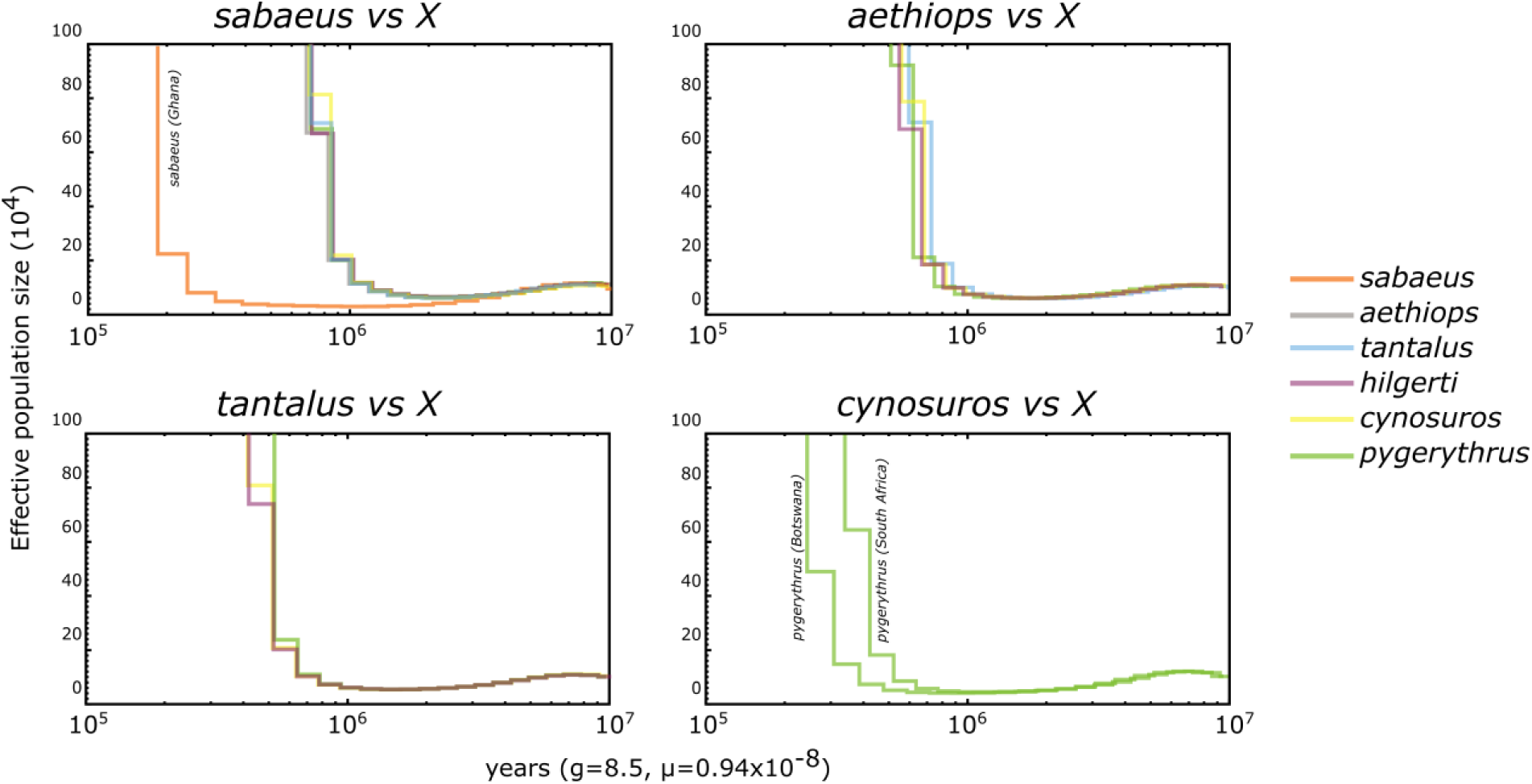
Artificial hybrid PSMC (hPSMC) within the vervet clade. Same as Figure S4, estimating divergence-times within the vervet genus.

**Figure S7.**
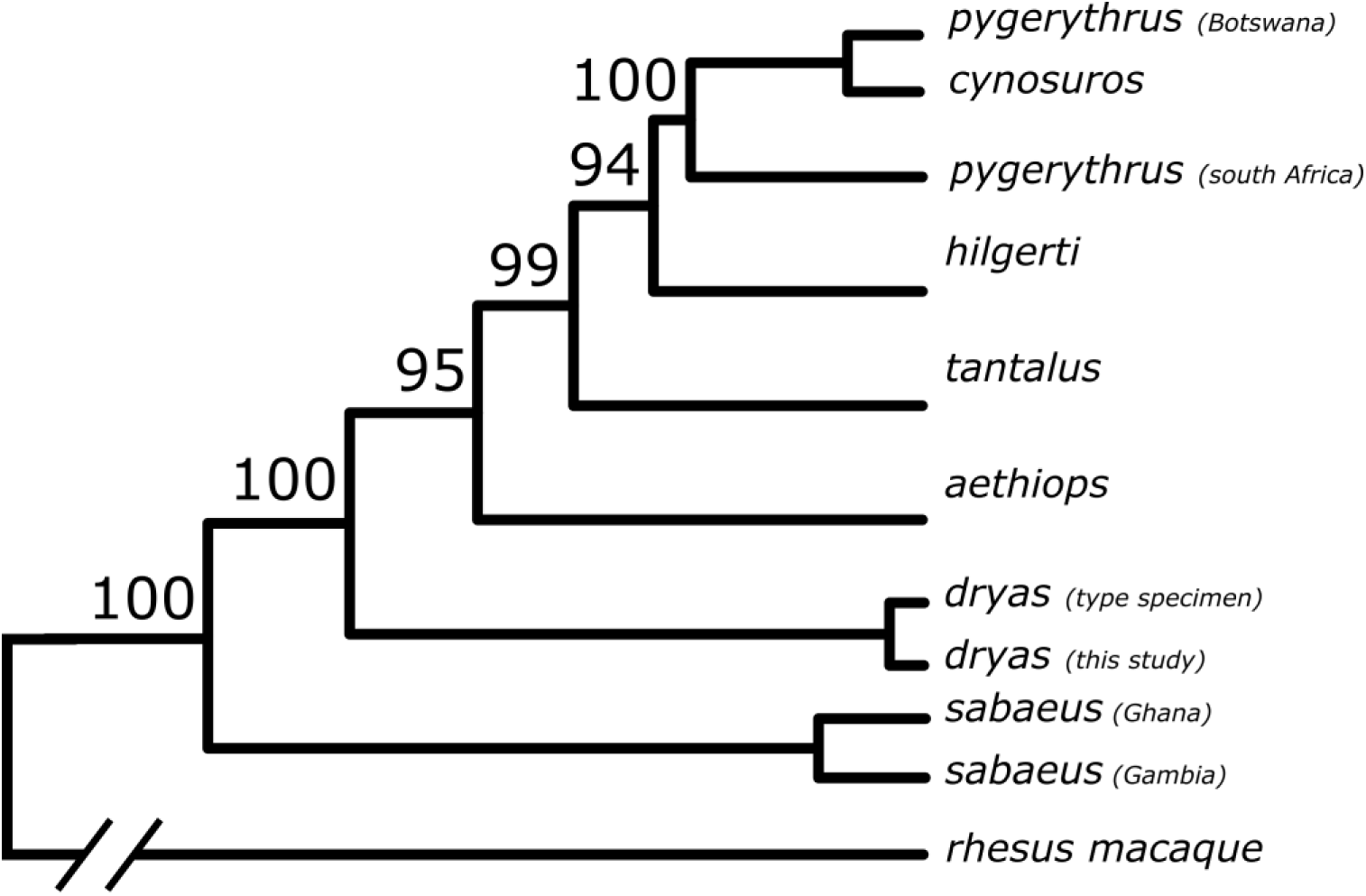
Maximum likelihood tree based on complete mitochondrial genomes for the vervets and the Dryas monkey (same as Figure 1C). Numbers depict bootstrap support values (1000 bootstrap replicates were run).

**Figure S8.**
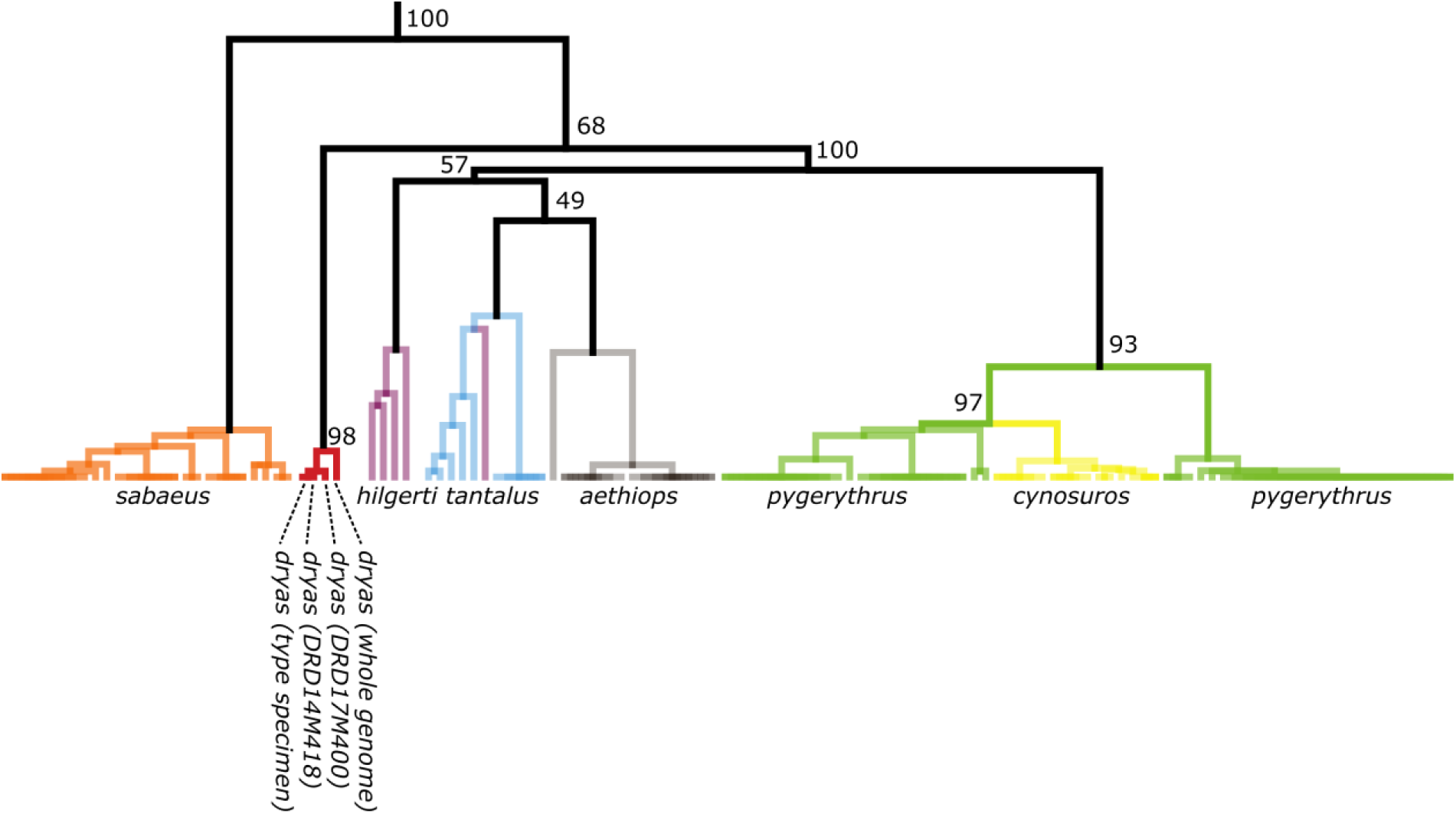
Maximum likelihood tree based on the cytochrome B sequence for the vervets and the Dryas monkey. Numbers depict bootstrap support values (1000 bootstrap replicates were run). Note the close clustering of all available Dryas monkey cytochrome B sequences.

**Figure S9.**
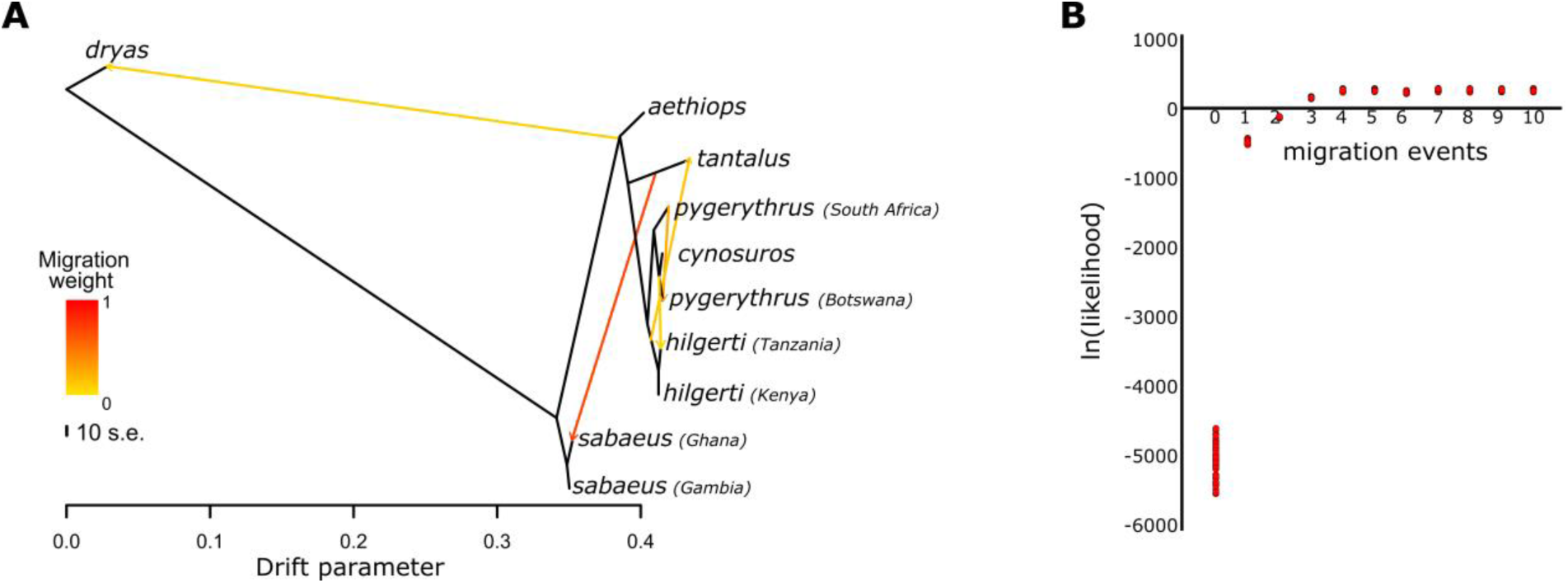
Model-based estimates of gene-flow. **(A)** TreeMix analysis for five migration events (m=5). Population allele frequencies are separated by species and geographic origin. **(B)** Likelihood support values for TreeMix models with 0 to 10 migration events respectively. After modelling five migration events, the model likelihood does not increase any further.

**Figure S10.**
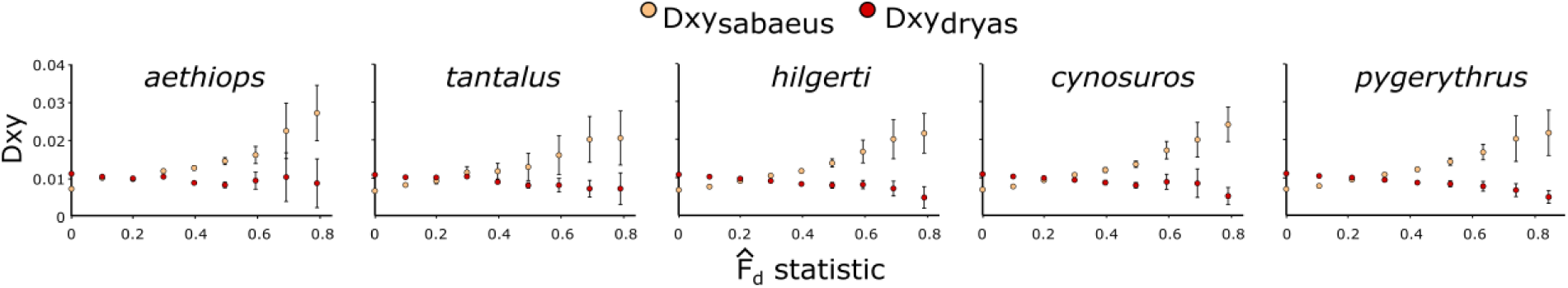
Identifying introgressed regions with Dxy statistic stratified by 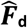. Errors bars show ±3SE. Window size = 10kb. Windows with high 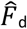 statistic (the excess of shared derived alleles with the Dryas monkey over that with *Chl. sabaeus*) have on average unusual large genetic distance (D_xy_) to *Chl. sabaeus* and unusual low genetic distance to the Dryas monkey. Note that the vast majority of windows is at 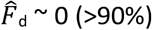. D_xy-sabaeus_ at windows with 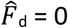 (putative non-introgressed windows) is generally around 0.007, whereas D_xy-dryas_ at these windows ∼0.012. This is in agreement with the obtained divergence time estimates (Figure 1B) (e.g. genetic distance of the non-*sabaeus* vervets to the Dryas monkey is on average 1.75 - 2.25 times larger than to *Chl. sabaeus*). At high 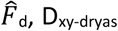 is around ½-⅓ that of 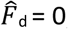, suggesting that these windows diverged on average half to one third as long ago as the other windows (thus ∼750.000 to 430.000 years ago as the most likely time period of introgression, if we assume the divergence of the Dryas monkey to the vervets to be ∼1.4 Mya, as indicated by our analyses).

**Figure S11.**
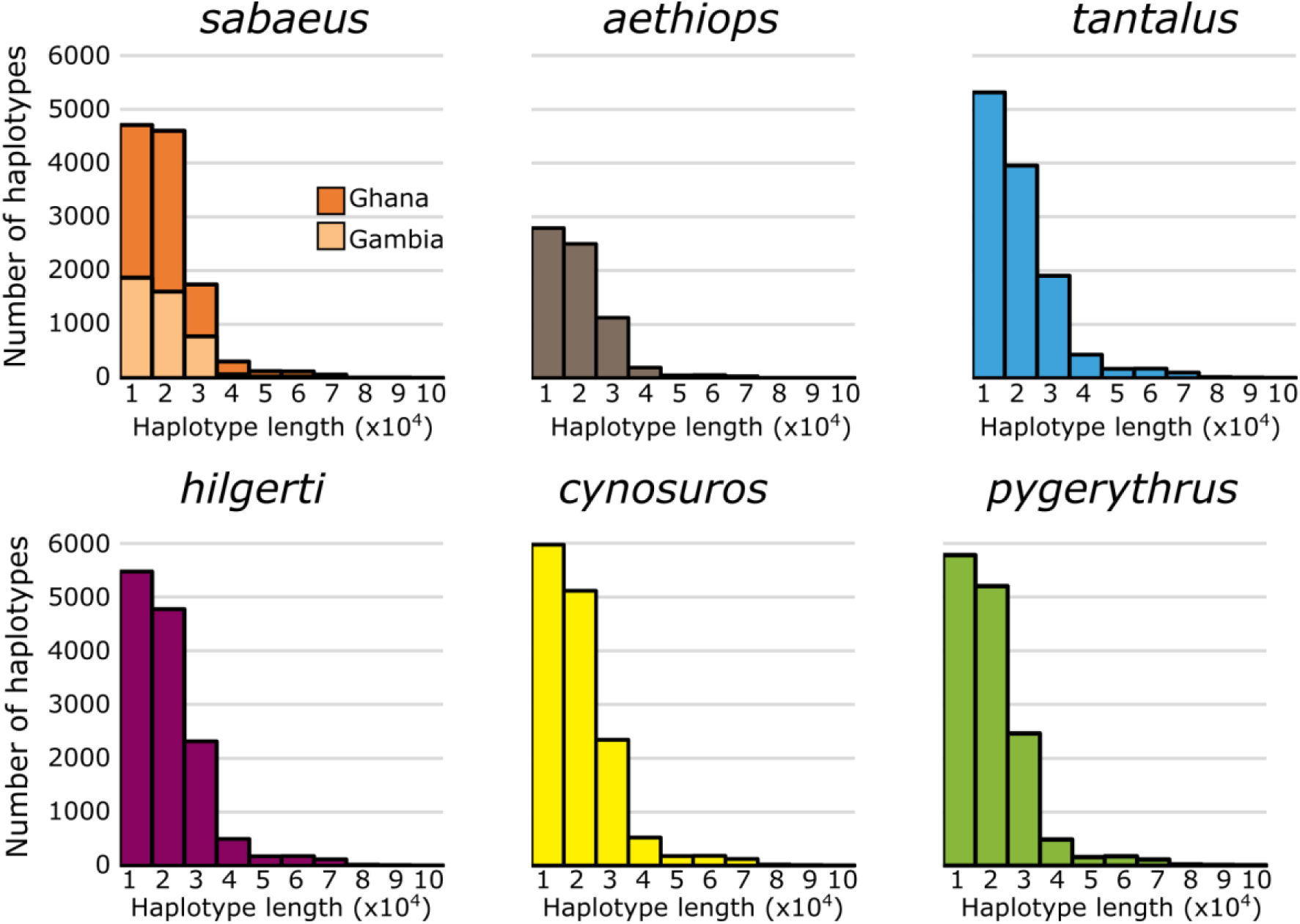
Length distribution of putatively introgressed haplotypes. Windows are short on average for all species. *Chl. sabaeus* individuals from Gambia have the lowest number of putatively introgressed windows, which were likely introduced into this species via secondary gene-flow with *Chl. tantalus*. Ghana individuals have a much higher number of these windows, as a result of the recent strong secondary admixture with *Chl. tantalus*.

**Figure S12.**
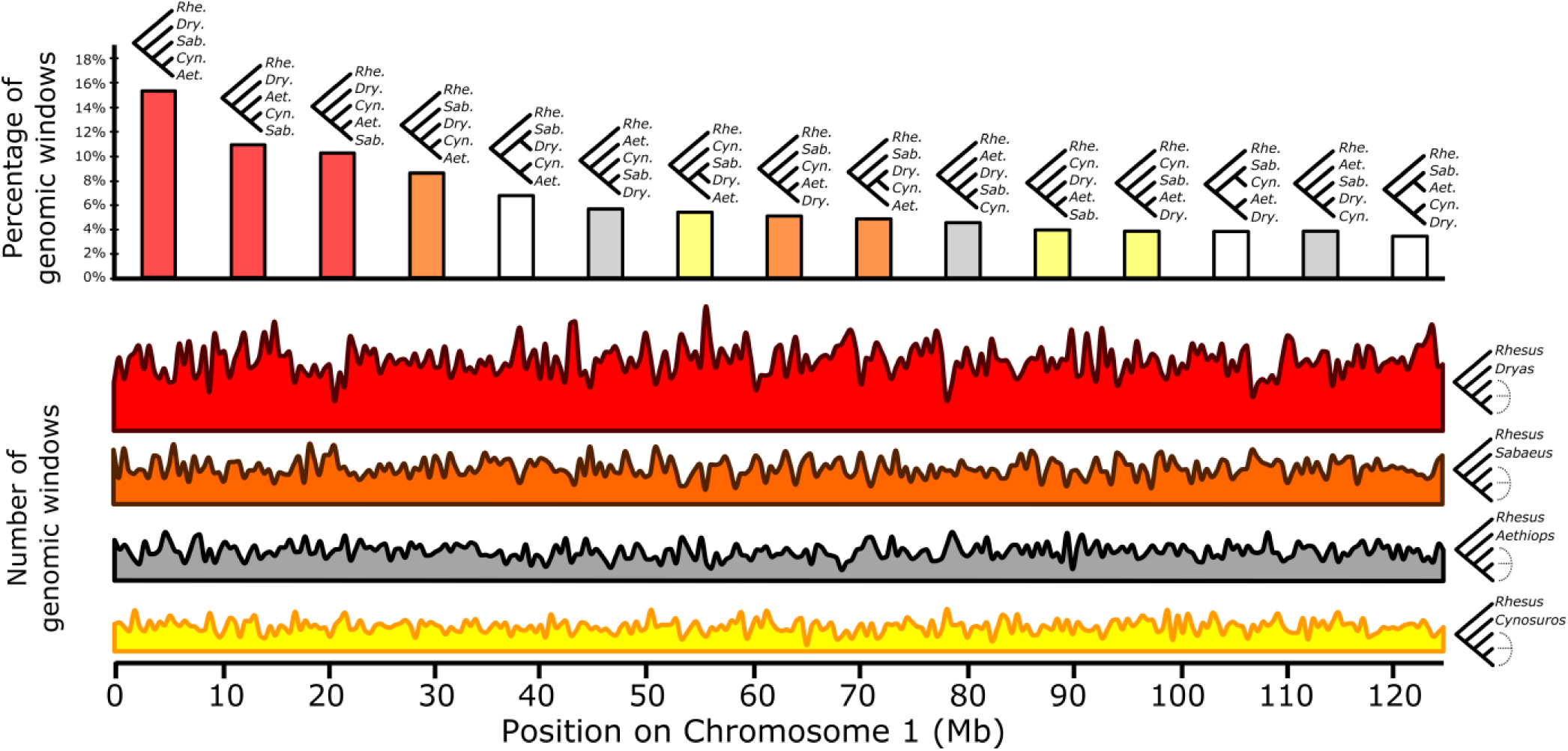
Exploring evolutionary relationships across the genome using topology weighting (*Twisst*). The top graph shows how often a depicted topology is supported among all non-overlapping 50 SNP windows (Rhe. = Rhesus macaque, Dry. = Dryas monkey, Sab. = *Chl. sabaeus*, Cyn. = *Chl. cynosuros*, Aet. = *Chl. aethiops*). The vast majority of windows supports the global topology (red-bars), in which the Dryas monkey is sister to all vervets. A smaller fraction of windows supports different topologies, likely to be the result of introgression and incomplete lineage sorting. Note that the second-most supported topology places the Dryas monkey inside the vervet clade with *Chl. sabaeus* as sister to all (orange bars). Plotting the occurrence of the different topologies along the genome (below), we did not identify long genomic regions supporting a contrasting topology to the majority topology (only chromosome 1 is shown here).

